# Macrophage-produced peroxynitrite induces antibiotic tolerance and supersedes intrinsic mechanisms of persister formation

**DOI:** 10.1101/2021.05.20.445079

**Authors:** Jenna E. Beam, Nikki J. Wagner, John C. Shook, Edward S.M. Bahnson, Vance G. Fowler, Sarah E. Rowe, Brian P. Conlon

**Affiliations:** Department of Microbiology and Immunology, University of North Carolina-Chapel Hill, Chapel Hill, North Carolina 27599, USA; Marsico Lung Institute, University of North Carolina at Chapel Hill, Chapel Hill, North Carolina 27599, USA; Department of Surgery, Division of Vascular Surgery, University of North Carolina at Chapel Hill, North Carolina 27599, USA; Center for Nanotechnology in Drug Delivery, University of North Carolina at Chapel Hill, NC; Curriculum in Toxicology & Environmental Medicine, University of North Carolina at Chapel Hill, North Carolina 27599, USA; McAllister Heart Institute, University of North Carolina at Chapel Hill, North Carolina 27599, USA; Department of Cell Biology & Physiology. University of North Carolina at Chapel Hill, North Carolina 27599, USA; Division of Infectious Diseases, Duke University School of Medicine, Durham, North Carolina 27710, USA

## Abstract

*Staphylococcus aureus* is a leading human pathogen that frequently causes chronic and relapsing infections. Antibiotic tolerant persister cells contribute to frequent antibiotic failure in patients. Macrophages represent an important niche during *S. aureus* bacteremia and recent work has identified a role for oxidative burst in the formation of antibiotic tolerant *S. aureus*. We find that host-derived peroxynitrite, the reaction product of superoxide and nitric oxide, is the main mediator of antibiotic tolerance in macrophages. Using a collection of *S. aureus* clinical isolates, we find that, despite significant variation in persister formation in pure culture, all strains were similarly enriched for antibiotic tolerance following internalization by activated macrophages. Our findings suggest that host interaction strongly induces antibiotic tolerance and may negate bacterial mechanisms of persister formation, established in pure culture. These findings emphasize the importance of studying antibiotic tolerance in the context of bacterial interaction with the host suggest that modulation of the host response may represent a viable therapeutic strategy to sensitize *S. aureus* to antibiotics.

## INTRODUCTION

*Staphylococcus aureus* is a Gram-positive bacterial pathogen that frequently causes chronic and relapsing infections, ranging from relatively minor skin and soft tissue infections to more serious infections like necrotizing pneumonia and bacterial sepsis (1). Despite the availability of antibiotics to treat *S. aureus* infections, treatment failure is common (2, 3). Antibiotic tolerance is the capacity of bacterial cells to survive for extended periods in the presence of a bactericidal antibiotic and subpopulations of antibiotic tolerant cells, called persisters, are frequently implicated in antibiotic treatment failure (4-6). Despite the apparent clinical importance of antibiotic tolerance and persister cell formation, the mechanism(s) underlying their formation during infection remain unclear (7-9).

Antibiotic tolerance and persister cell formation have long been studied using pure bacterial cultures under in vitro conditions. Numerous mechanisms have been identified using this approach, including toxin-antitoxin modules, induction of the stringent response, reduced respiration, ATP depletion, and metabolic collapse (8, 10-17). However, the relative contribution of these bacterial mechanisms to antibiotic survival during host interaction remains unclear. Recent work from our lab and others has identified a role for the host response in inducing antibiotic tolerance (7, 8). Upon bacteremia, *S. aureus* is rapidly phagocytosed by macrophages and resides in the mature phagolysosome (18, 19). Although macrophages are effective at killing *S. aureus*, sub-populations of bacterial cells survive and these survivors are highly tolerant to antibiotics (7, 8, 19). We previously showed that macrophage-derived reactive oxygen species (ROS) collapse tricarboxylic acid (TCA) cycle activity and deplete ATP, leading to a metabolic state that is incompatible with antibiotic killing (7).

During oxidative burst, a myriad of reactive oxygen and nitrogen species (ROS/RNS) are produced (Fig 1A). The NADPH oxidase (NOX) complex located on the phagosomal membrane generates superoxide anions from the reduction of molecular oxygen. These superoxide anions are generated inside the phagosomal lumen and poorly penetrate the bacterial membrane (20). Superoxide can be dismutated into oxygen and hydrogen peroxide. The latter is further detoxified into oxygen and water. This occurs spontaneously, but *S. aureus* also produces a catalase enzyme that facilitates this detoxification. In addition to the production of superoxide, the inducible nitric oxide synthase (iNOS) protein located in the macrophage cytoplasm converts L-arginine to nitric oxide. Nitric oxide freely diffuses into the phagosomal lumen, as well as through the bacterial cell membrane (21). Nitric oxide can be detoxified by *S. aureus* through the Hmp and Nar proteins (22, 23). Additionally, superoxide and nitric oxide react at a diffusion-controlled rate to form the potent reactive species, peroxynitrite (24). Once formed, peroxynitrite can oxidize and nitrate various cellular components, including nucleic acids, proteins, and lipids (24-26). Importantly, the rate of formation of peroxynitrite from nitric oxide and superoxide is an order of magnitude faster than that of the superoxide dismutase-catalyzed conversion of superoxide into hydrogen peroxide, suggesting that peroxynitrite levels are likely highly abundant during oxidative burst, when superoxide and nitric oxide are both produced (24).

**Figure 1.**
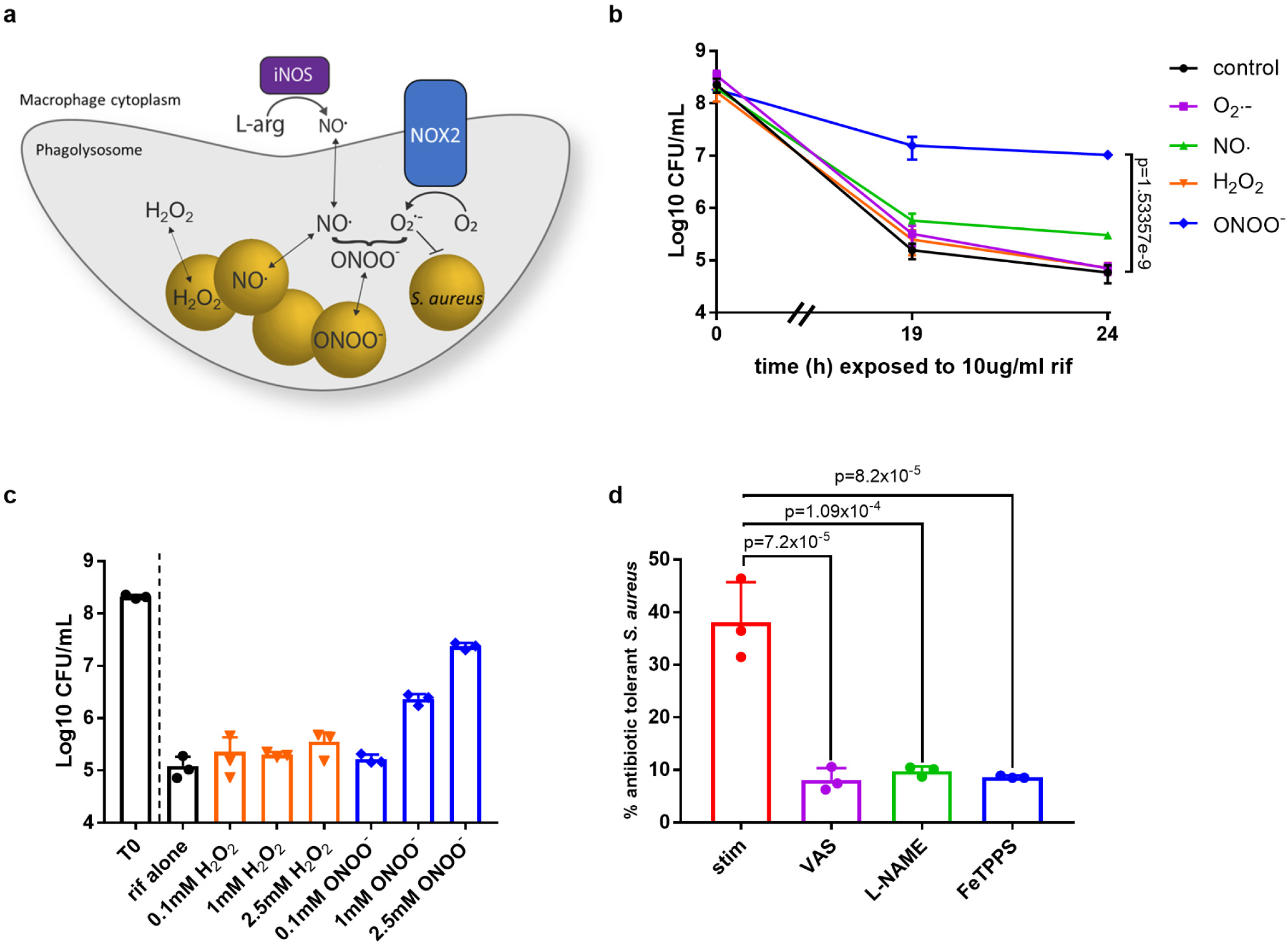
Peroxynitrite is the main driver of tolerance to rifampicin in macrophages. (a) Schematic representing the production of reactive oxygen and nitrogen species produced during oxidative burst in macrophages. (b) *S. aureus* strain HG003 was grown to mid-exponential phase in vitro and exposed to 0.3U/L xanthine oxidase (O_2_·-), 1mM DEA NO/NOate (NO·), 2.5mM H_2_O_2_, or 2.5 mM peroxynitrite (ONOO-) for 20min prior to the addition of rifampicin at time 0. At indicated times, an aliquot was removed, washed in 1% NaCl and plated to enumerate survivors. (c) *S. aureus* strain HG003 was grown to mid-exponential phase in vitro and exposed to 0.1mM, 1mM, or 2.5mM peroxynitrite (blue bars) or H_2_O_2_ (orange bars) for 20min prior to the addition of rifampicin (10μg/mL) at time 0. At indicated times, an aliquot was removed, washed in 1% NaCl and plated to enumerate survivors. (d) % survival of *S. aureus* cells after internalization by J774 macrophages that were stimulated overnight with LPS/IFNγ, then treated with either 10μM VAS, 2mM L-NAME, or 200µM FeTPPS for 1h prior to infection. 10μg/ml rifampicin (rif) was added at time 0, after 30min incubation with *S. aureus* to allow for phagocytosis. After 4h rifampicin treatment, *S. aureus* survival was compared to the untreated control. (b-d) Data are representative of n=3 biological samples and all experiments were repeated 3 times on separate days. Error bars represent standard deviation. Statistical significance was determined using One-Way ANOVA with Sidak’s multiple comparison test.

In the current study, we aimed to identify which reactive species was predominantly responsible for the induction of antibiotic tolerance in phagocytosed *S. aureus*. We found that peroxynitrite generated by activated macrophages indiscriminately induced antibiotic tolerance in a variety of *S. aureus* strains, including clinical bacteremia isolates and mutants that displayed decreased persister formation in vitro. These findings suggest that targeting macrophage-produced peroxynitrite may improve antibiotic efficacy against recalcitrant *S. aureus* populations in the host and that antibiotic tolerance needs to be investigated in the context of host interaction if physiologically relevant mechanisms of tolerance are to be overcome.

## RESULTS

### Macrophage-derived peroxynitrite induces tolerance to rifampicin

We previously reported that ROS produced by stimulated macrophages induced antibiotic tolerance of internalized *S. aureus* cells (7). To determine which reactive species is the main driver of intracellular antibiotic tolerance, we tested the capacity of superoxide, nitric oxide, peroxynitrite, and hydrogen peroxide to induce rifampicin tolerance in vitro. *S. aureus* was grown to exponential phase and exposed to various sources of reactive species for 20 minutes prior to rifampicin challenge. Redox cycling agents such as menadione and paraquat have been used extensively to induce superoxide production (7). However, these agents generate superoxide within the bacterial cytoplasm, which does not accurately recapitulate the production of superoxide in the phagosomal lumen, which poorly penetrates the bacterial membrane (20). To better replicate extracellular superoxide production in vitro, *S. aureus* was exposed to xanthine oxidase (XO) with acetaldehyde, a well-characterized enzyme that produces superoxide in the growth media (27), prior to antibiotic challenge. Neither superoxide nor nitric oxide, produced via the nitric oxide donor DEA NONOate, induced tolerance to rifampicin (Fig 1B). In contrast, pre-treatment with peroxynitrite induced complete tolerance to rifampicin. Hydrogen peroxide was previously shown to induce tolerance when added at high levels (120mM)(7). However, at equivalent concentrations to peroxynitrite, hydrogen peroxide had no impact on rifampicin tolerance (Fig 1B). Addition of these oxidizing agents in the absence of rifampicin did not cause bacterial cell death (S1A Fig). We then examined if the induction of antibiotic tolerance by peroxynitrite was concentration dependent by challenging cultures with increasing concentrations prior to rifampicin treatment. At sub-lethal peroxynitrite concentrations (S1B Fig), tolerance to rifampicin was induced in a dose-dependent manner (Fig 1C). Equivalent concentrations of hydrogen peroxide had no impact on antibiotic susceptibility (Fig 1C, S1B Fig). These results suggest that although superoxide, hydrogen peroxide or nitric oxide may have the capacity to induce antibiotic tolerance, peroxynitrite does so with far greater potency.

Next, to assess the relative contribution of each species during macrophage infection, J774 macrophages were treated with inhibitors of oxidative and nitrosative burst prior to infection with *S. aureus* and subsequent exposure to rifampicin. Macrophages were treated with inhibitors of burst for 1h, followed by infection with *S. aureus* for 30 minutes before the addition of rifampicin to allow for bacterial uptake in the absence of antibiotic. Addition of VAS2870 (VAS), an inhibitor of the NOX complex (which produces superoxide)(28), to the J774 macrophages restored rifampicin susceptibility of intracellular *S. aureus* (Fig 1D, S1C Fig). Similarly, addition of L-NAME, which inhibits inducible nitric oxide synthase (iNOS; which produces nitric oxide)(29), to the J774 macrophages restored rifampicin susceptibility of intracellular *S. aureus* (Fig 1D, S1C Fig). Since superoxide and nitric oxide spontaneously react to produce peroxynitrite, we hypothesized that peroxynitrite was driving antibiotic tolerance of intracellular *S. aureus*. In support of this, addition of FeTPPS, a peroxynitrite decomposition catalyst (30), also sensitized *S. aureus* to rifampicin (Fig 1D, S1C Fig). Taken together, these data suggest that peroxynitrite is primarily responsible for the induction of antibiotic tolerance in phagocytosed *S. aureus*.

Because the duration of oxidative burst is limited and high levels of ROS are only transiently produced (31, 32), we examined if the tolerant state was maintained after 2 hours of internalization (compared to 30 minutes in previous experiments). Indeed, we found tolerance was stable for at least that duration, further suggesting the physiological relevance of the phenotype (S1D Fig).

### Exposure to peroxynitrite reduces TCA cycle activity through post-translational modification of aconitase

We have previously shown that exposure to ROS, through treatment with menadione, reduces TCA cycle activity and depletes ATP levels in *S. aureus*, thus conferring protection from antibiotic killing (7). Since peroxynitrite appears to be the main mediator of antibiotic tolerance of *S. aureus* internalized by macrophages, we aimed to determine if peroxynitrite does so by through TCA cycle inactivation. We found that only *S. aureus* exposed to peroxynitrite displayed reduced levels of intracellular ATP (Fig 2A), while *S. aureus* exposed to extracellular superoxide, nitric oxide, or hydrogen peroxide had similar levels of ATP to control untreated cells (Fig 2A). ATP levels were also significantly reduced in *S. aureus* recovered from stimulated macrophages compared to unstimulated macrophages (Fig 2B).

**Figure 2.**
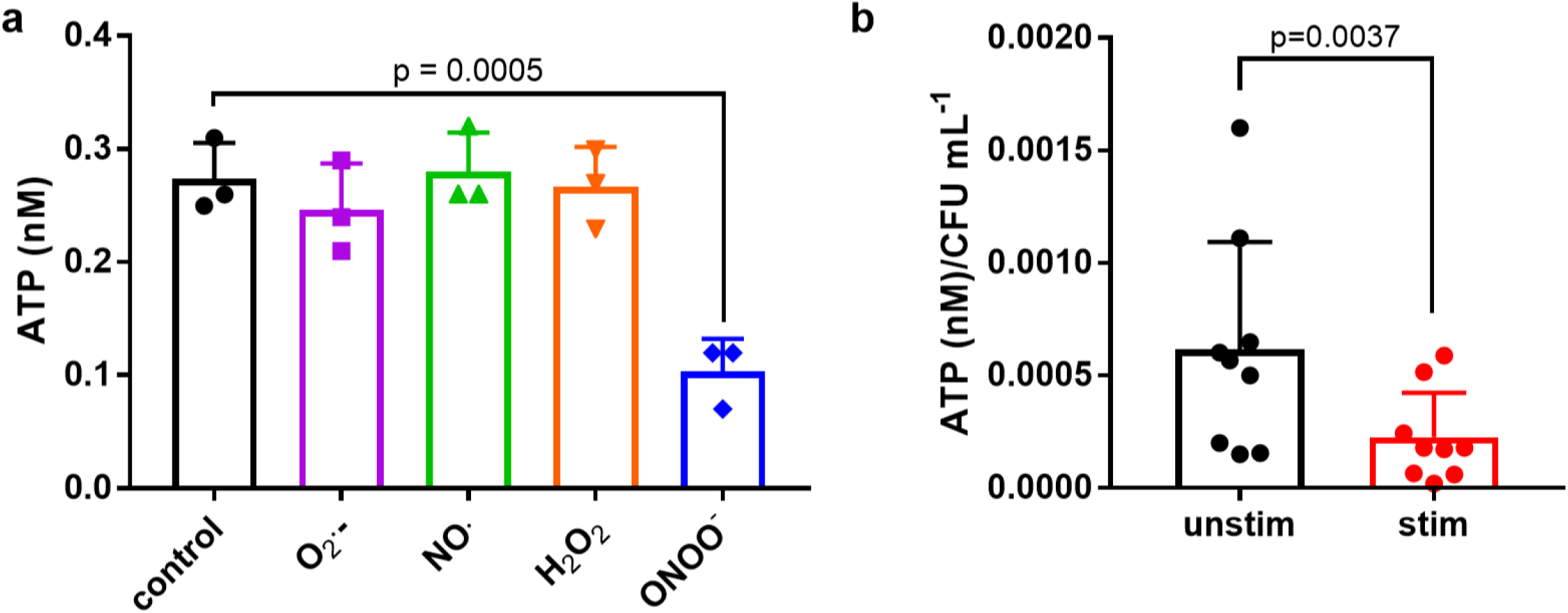
ATP depletion correlates with antibiotic tolerance. (a) *S. aureus* strain HG003 was grown to the mid-exponential phase and treated with 0.3U/L xanthine oxidase (O_2_·-), 1mM DEA NO/NOate (NO·), 2.5mM H_2_O_2_, or 2.5 mM peroxynitrite (ONOO-) for 30 min. Intracellular ATP was measured using a BacTiter-Glo cell viability assay. (b) Intracellular ATP was measured using a BacTiter-Glo cell viability assay in *S. aureus* recovered from unstimulated and stimulated macrophages. Macrophages were infected at MOI 50 with *S. aureus* strain HG003 for 0.5h. (a) Data are representative of n=3 biological samples and all experiments were repeated 3 times on separate days. (b) n=9 biologically independent samples. Error bars represent the s.d. Statistical significance was determined using a one-way analysis of variance (ANOVA) with Sidak’s multiple comparison test (a) or unpaired two-tailed Student’s t-test (b).

Aconitase is an iron-sulfur (Fe-S) cluster containing enzyme known to be extremely sensitive to oxidative stress (33). Peroxynitrite has previously been shown to damage and inactivate mammalian aconitase (33). To confirm that *S. aureus* aconitase is sensitive to peroxynitrite, we exposed exponential phase cultures to peroxynitrite and measured aconitase activity. Exposure to peroxynitrite reduced aconitase activity in vitro (Fig 3A, S2A Fig). Using a *S. aureus* aconitase mutant strain, our prior work indicates a critical role for aconitase in *S. aureus* antibiotic tolerance (7). Here, we demonstrate that a *S. aureus* aconitase mutant exhibits increased tolerance to rifampicin relative to wildtype in vitro in the absence of ROS, suggesting that aconitase inactivation is sufficient to induce antibiotic tolerance (Fig 3B).

**Figure 3.**
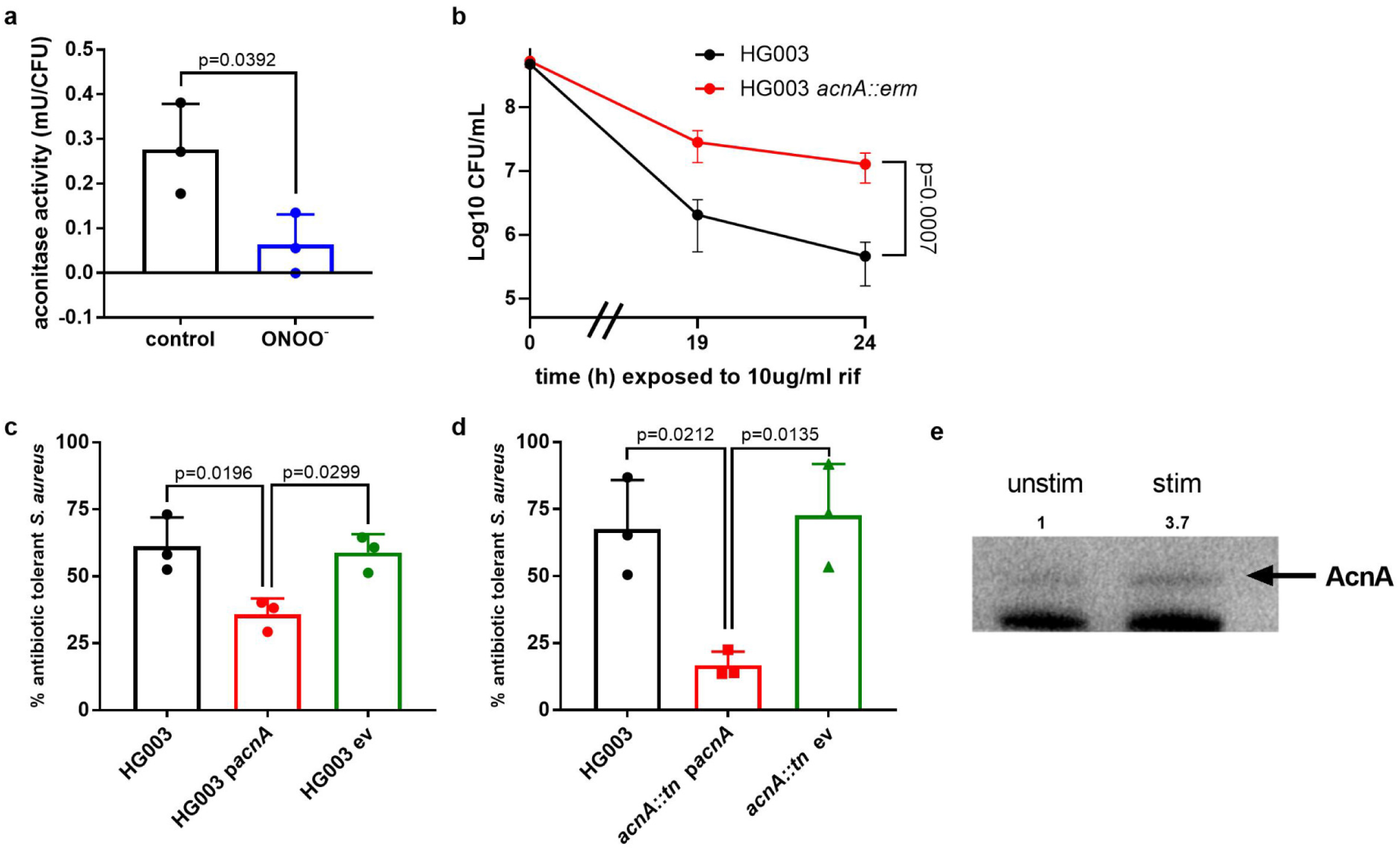
Peroxynitrite-mediated aconitase inactivation is the main driver of antibiotic tolerance. (a) *S. aureus* strain HG003 was grown to mid-exponential phase and treated with 2.5mM peroxynitrite for 30min, and aconitase activity was measured without activation according to manufacturer’s instructions. The averages of n = 3 biologically independent samples are shown. (b) Survival of *S. aureus* strain HG003 (black) or HG003 *acnA::erm* (red). Cells were grown to mid-exponential phase, then treated with 10μg/ml rifampicin (rif). At indicated times, an aliquot was removed, washed in 1% NaCl and plated to enumerate survivors. (c) % survival of *S. aureus* strain HG003 (black), HG003 pEPSA5_*acnA*_FLAG (*acnA* overexpression; red), and HG003 pEPSA5 empty vector (green) after internalization by J774 macrophages that were stimulated overnight with LPS + IFNγ. 10μg/ml rifampicin (rif) was added at time 0, after 30min incubation with *S. aureus* to allow for phagocytosis. Prior to macrophage infection, all strains were induced with 2% xylose for 1h. (d) % survival of *S. aureus* strain HG003 (black), HG003 *acnA::ermB* pEPSA5_*acnA*_FLAG (*acnA* overexpression; red), and HG003 *acnA::ermB* pEPSA5 empty vector (green) after internalization by J774 macrophages that were stimulated overnight with LPS + IFNγ. 10μg/ml rifampicin (rif) was added at time 0, after 30min incubation with *S. aureus* to allow for phagocytosis. Prior to macrophage infection, all strains were induced with 0.2% xylose for 1h. (e) Western blot for 3-nitrotyrosine (YNO_2_). *S. aureus* strain HG003 *acnA::ermB* pEPSA5_*acnA_*FLAG was induced with 0.2% xylose for 1h. Unstimulated (unstim) and stimulated (stim) macrophages were infected for 4h. *S. aureus* cells were collected and *S. aureus* AcnA was immunoprecipitated using anti-FLAG magnetic beads and blotted for YNO_2_. Non-specific band used as loading control. (a-d) Data are representative of n=3 biological samples and all experiments were repeated 3 times on separate days. In (a-d) error bars represent the s.d. Statistical significance was determined using a one-way analysis of variance (ANOVA) with Sidak’s multiple comparison test (b-d) or unpaired two-tailed Student’s t-test (a). (d) Blot is representative of n=3 IP-Western blot analyses performed on separate days. Bands were quantified using densitometry (FIJI).

If aconitase inactivation is the main driver of antibiotic tolerance, we reasoned that overexpression of aconitase would restore susceptibility in stimulated macrophages. To test this, J774 macrophages were infected with a *S. aureus* strain overexpressing aconitase (34). Overexpression of aconitase in *S. aureus* during macrophage infection was sufficient to sensitize *S. aureus* to antibiotic killing (Fig 3C-D, S2B Fig and S2C Fig), indicating that host-mediated inactivation of aconitase is the primary mediator of tolerance in macrophages.

Next, we aimed to directly measure peroxynitrite-mediated damage of aconitase. Nitration of tyrosine residues is an established method that can be used as a fingerprint of peroxynitrite-mediated damage of proteins (35, 36). It should be noted that studies using purified mitochondrial aconitase suggest that peroxynitrite-mediated disruption of the [4Fe-4S]^2+^ cluster of aconitase is most likely responsible for enzyme inactivation, although nitration of tyrosine residues and oxidation of cysteine residues may contribute to activity loss (33, 37-39). We infected J774 macrophages with *S. aureus* expressing a FLAG-tagged aconitase (34). Following infection, bacterial aconitase was purified by immunoprecipitation (IP) of the FLAG-tag, followed by Western blot for 3-nitrotyrosine (3-NT; Fig 3E). Compared to unstimulated macrophages (correlating to decreased ROS production), *S. aureus* aconitase purified from stimulated macrophages exhibited increased levels of 3-NT (Fig 3E). Taken together, these data suggest that peroxynitrite-mediated damage of *S. aureus* aconitase is sufficient to induce rifampicin tolerance.

### Peroxynitrite induces antibiotic tolerance in *S. aureus* clinical bacteremia isolates

Using a collection of *S. aureus* clinical isolates from bacteremia patients, we identified major variability in antibiotic tolerance under in vitro conditions, despite the isolates exhibiting similar minimum inhibitory concentrations to rifampicin (Fig 4A, S3A Fig). Using 3 low persister clinical isolates (BC1263, BC1266, and BC1272) and 3 high persister clinical isolates (BC1267, BC1271, and BC1274) identified in an in vitro screen (Fig 4A), we examined antibiotic tolerance following phagocytosis by stimulated or unstimulated macrophages. Strikingly, all bacteremia isolates were highly induced for rifampicin tolerance in stimulated macrophages and multiple log-scale differences in persister formation identified in vitro were no longer evident (Fig 4B, S3B-G Fig).

**Figure 4.**
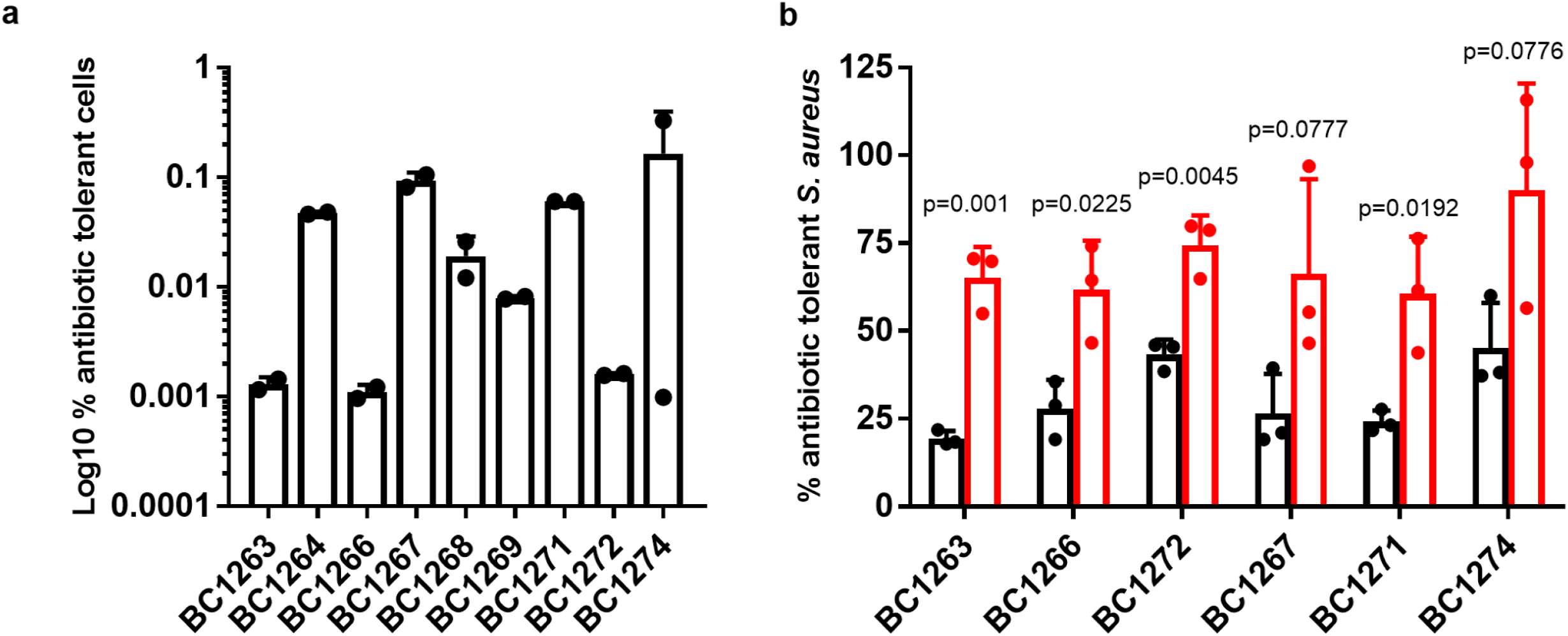
Antibiotic tolerance is driven by host cell interactions. (a) % survival of *S. aureus* bacteremia clinical isolates following 24h rifampicin treatment. (b) % survival of *S. aureus* bacteremia clinical isolates after internalization by unstimulated (black) J774 macrophages or J774 macrophages stimulated overnight with LPS/IFNγ (red). 10μg/ml rifampicin (rif) was added at time 0, after 30min incubation with *S. aureus* to allow for phagocytosis. *S. aureus* was treated with rifampicin for 4h compared to the untreated control. (a) Data are representative of n=2 biological samples and all experiments were repeated 3 times on separate days. (b) Data are representative of n=3 biological samples and all experiments were repeated 3 times on separate days. (b) Error bars represent the s.d. Statistical significance was determined using unpaired two-tailed Student’s t-test.

While numerous mechanisms of antibiotic tolerance and persister formation have been established in vitro, we were interested in determining the relative importance of these mechanisms in the survival of phagocytosed *S. aureus* to antibiotics. Activation of the stringent response (SR) was shown to contribute to the formation of intracellular *S. aureus* persisters in unstimulated macrophages (8) and TA modules were shown to contribute to antibiotic tolerance in *S. aureus* biofilms (10). We infected unstimulated and stimulated J774 macrophages with a SR-deficient RSH synthase-domain mutant (*Δrsh*_*syn*_(40)) and its wildtype HG001 control, as well as a triple TA module mutant (Δ3TA(12)) and its wildtype Newman control. Interestingly, stimulated macrophages significantly induced tolerance to rifampicin in both the *Δrsh*_*syn*_ and Δ3TA mutants (S4 Fig). Stimulated macrophages also induced tolerance to moxifloxacin in the *Δrsh*_*syn*_ mutant similarly to wildtype (S5E Fig). Together, these data indicate that host-derived ROS-mediated tolerance is the dominant driver of antibiotic tolerance within stimulated macrophages.

## DISCUSSION

Within minutes of entering the blood, macrophages engulf *S. aureus* (19). The initial pro-inflammatory macrophage response is critical for preventing *S. aureus* infection. However, if the infection establishes, we find that high levels of ROS can induce antibiotic tolerance, which may contribute to treatment failure. Specifically, we identify peroxynitrite, the reaction product of superoxide and nitric oxide, to be the main driver of ROS-mediated antibiotic tolerance. Peroxynitrite is a potent reactive species capable of nitrating and oxidizing various cellular components, notably Fe-S cluster containing enzymes (24, 35). We find that peroxynitrite-mediated damage of *S. aureus* aconitase leads to collapse of the TCA cycle, reduction of ATP, and ultimately entrance into a metabolic state that is incompatible with antibiotic killing. In support of our findings, Huemer *et al*., recently demonstrated that antibiotic tolerant *S. aureus* recovered from a patient abscess displayed reduced levels of ATP and aconitase activity, despite increased transcription and cytosolic accumulation of aconitase (9).

Why peroxynitrite drives the formation of antibiotic tolerant *S. aureus* in macrophages over other reactive species (ie hydrogen peroxide) may be attributed to multiple factors. Firstly, the formation of peroxynitrite from superoxide and nitric oxide is extremely efficient, occurring an order of magnitude faster than that of the conversion of superoxide into hydrogen peroxide, suggesting levels of peroxynitrite are likely extremely high during oxidative burst (24). Secondly, since the phagolysosome is highly acidic, peroxynitrite will be rapidly protonated and thus able to freely pass through the bacterial membrane (24). Additionally, aconitase is characteristically highly sensitive to oxidative damage and the ability of peroxynitrite to efficiently cause aconitase damage and TCA cycle collapse in mammals is well-documented (33, 37-39, 41). Aside from damage to aconitase, additional non-lethal bacterial cell modifications induced by peroxynitrite may have importance in host-driven antibiotic tolerance in *S. aureus* as peroxynitrite reacts with thiols and sulfur-containing moieties three orders of magnitude faster than that of hydrogen peroxide (42, 43).

Scavenging peroxynitrite exogenously or repairing peroxynitrite-mediated damage have been shown to improve disease outcome in a variety of diseases, including ischemic stroke and septic shock (44-46). There are both direct and indirect methods for resolving peroxynitrite-mediated damage. Thiols, like glutathione and bacillithiol, directly scavenge peroxynitrite via reduction reaction, whereas ascorbic and uric acid indirectly resolve damage by inhibiting tyrosine nitration (47, 48). Metallic porphyrins, like FeTPPS and WW-85, facilitate decomposition of peroxynitrite by catalyzing the isomerization of peroxynitrite into nitrate (44, 46, 49). WW-85 has been shown to diminish the need for arginine vasopression treatment during methicillin-resistant *S. aureus* (MRSA) septic shock, improving disease outcome (46).

Historically, mechanisms underlying antibiotic tolerant and persister cell formation have been studied under in vitro conditions using pure bacterial cultures. Although numerous mechanisms contributing to persister formation have been identified this way, whether or not these mechanisms contribute to antibiotic tolerance in stimulated macrophages is unknown. We find that previously identified intrinsic bacterial mechanisms underlying persister formation are negated by the powerful tolerance inducing effects of macrophage-derived ROS. Further, using a collection of clinical isolates, we find that despite variability in persister formation in vitro, all bacteremia strains examined were highly tolerant in stimulated macrophages. Altogether, these data suggest that host-derived peroxynitrite is capable of indiscriminately inducing antibiotic tolerance in an array of genetically diverse *S. aureus* isolates.

Overall, our results identify host-derived peroxynitrite as a major contributor to antibiotic tolerance during *S. aureus* infection. These findings suggest that acute modulation of peroxynitrite via inhibitors of peroxynitrite formation or peroxynitrite decomposition catalysts may represent a viable therapeutic strategy for improving antibiotic efficacy against recalcitrant *S. aureus* infection.

## MATERIALS AND METHODS

### Bacterial strains and growth conditions

*S. aureus* strains HG003, HG003 *acnA*::*erm*(7), Newman, Newman*ΔmazEFΔaxe1-txe1Δaxe2-txe2* (abbreviated Δ3TA)(12), HG001, and HG001Δ*rsh*_*syn*_(40) were routinely cultured in Mueller Hinton broth (MHB) at 37 °C and 225 r.p.m. Strains harboring plasmids pEPSA5 empty vector or pEPSA5::acnA_FLAG (34) were grown in the presence of 10µg/ml chloramphenicol and induced with 0.2% or 2% xylose where indicated. *S. aureus* bacteremia isolates were obtained under an IRB exemption from a pre-existing collection. Bacteremia isolates were cultures in MHB at 37 °C and 225 r.p.m.

### Macrophage growth and infection

Macrophage growth and infection was performed as described previously(7). Briefly, J774A.1 murine macrophage-like cells (ATCC TIB-67) were cultured in Dulbecco’s Modified Essential Media, high glucose (DMEM) (Gibco) supplemented with 10% fetal bovine serum (FBS) (Milipore), non-essential amino acids (NEAA) (Gibco), sodium pyruvate (Gibco) and L-glutamine (Gibco) in a humidified incubator at 37°C and 5% CO_2_. For infections, macrophages were seeded at a density of 2⨯10^5^ per well in 24 well plates in Minimum Essential Media (MEM) (Gibco) supplemented with 10% FBS and L-glutamine. For ATP assays, macrophages were seeded at 4⨯10^5^ cells/ml in MEM in 6 well plates. For IP-WB, macrophages were seeded at 6.5⨯105 cells/ml in MEM in 10cm tissue culture-treated Petri dishes. To stimulate the macrophages, 500ng/ml lipopolysaccharide (LPS) from *Escherichia coli* O55:B5 (Sigma) and 20ng/ml recombinant murine interferon-g (Peprotech) (IFNg) were added to supplemented MEM overnight(50).

*S. aureus* strains HG003, HG001, and Newman wild type or mutant strains were used to infect macrophages at a multiplicity of infection (MOI) of 50. Where indicated, macrophages were pre-treated with 10µM VAS2870 (VAS) (Cayman Chemicals), 2mM L-NG-Nitroarginine Methyl Ester (L-NAME) (Santa Cruz Biotechnology), or 200μM Fe(III)5,10,15,20-tetrakis(4-sulfonatophenyl)porphyrinato chloride (FeTPPS)(Cayman Chemicals) for 1h prior to infection. Plates were centrifuged at 1000xg for 2min to bring bacteria into contact with the macrophages. After 30 minutes or 2h (where indicated), the cells were washed once with PBS and fresh media containing 30µg/ml gentamicin (Fisher) was added to kill extracellular bacteria (51) and 10µg/ml rifampicin (Fisher) or 3μg/ml (50X MIC) moxifloxacin (Fisher) was added to the appropriate wells. At indicated times, macrophages were lysed with 0.1% Triton X-100 to release the bacteria. PBS was added to each well, lysates were resuspended by pipetting, serially diluted in 1% NaCl and plated to enumerate surviving bacteria. % survival after rifampicin or moxifloxacin treatment was determined by comparing survivors after 4h antibiotic treatment to survivors of the corresponding untreated 4h timepoint. Averages and standard deviations of 3 biological replicates are shown (n=3). Cell lines were obtained from UNC Lineberger Comprehensive Cancer Center’s Tissue Culture Facility. We did not authenticate or test cells for mycoplasma contamination. Statistical significance was calculated using the Student’s t-test (unpaired, two-tailed) or One-Way ANOVA with Sidak’s multiple comparison test as described in the figure legends.

### ROS measurements

The luminescent probe L-012 (Wako Chemical Corporation) was used to measure ROS. J774 macrophages were seeded at 2 × 10^4^ cells per well in white tissue-culture-treated 96-well plates. Macrophages were treated with either LPS + IFNγ or IL-4 + IL-13 as described above. The cells were washed three times with PBS. L-012 was diluted to 150 µM in Hanks’ balanced salt solution (Gibco). Luminescence was read immediately using a Biotek Synergy H1 microplate reader. Data shown are representative of 3 independent assays of 4 biological replicates. Statistical significance was calculated using a one-way ANOVA with Sidak’s multiple comparison test.

### Antibiotic survival assays

*S. aureus* strain HG003 and clinical isolates were cultured aerobically in MHB (Oxoid) at 37°C with shaking at 225 rpm for ∼16h. Stationary cultures were diluted 1:100 in MHB and grown to mid-exponential phase. An aliquot was plated to enumerate CFU (time 0) before the addition of antibiotics. Where indicated, the culture was incubated with 0.3U/L xanthine oxidase (Calbiochem), 1mM DEA NO/NOate (Cayman Chemical), 2.5mM H_2_O_2_, or 2.5 mM peroxynitrite (Cayman Chemical) for 20min prior to antibiotic challenge. For data in Fig. 1c, 0.1mM, 1mM, 2.5mM, 5mM, and 7mM H_2_O_2_ or peroxynitrite was used. Rifampicin was added at concentrations similar to the C_max_ in humans at recommended dosing (10µg/ml) (52). At indicated times, an aliquot was removed and washed with 1% NaCl. Cells were serially diluted and plated on tryptic soy agar (TSA) to enumerate survivors. Averages and standard deviations of 3 biological replicates are shown (n=3). Statistical significance was calculated using the Student’s t-test (unpaired, two-tailed) or One-Way ANOVA with Sidak’s multiple comparison test as described in the figure legends.

### Growth curves

*S. aureus* strain HG003 was cultured aerobically in Mueller-Hinton broth (MHB) (Oxoid) at 37°C with shaking at 225 rpm for ∼16h. Stationary cultures were diluted 1:100 in MHB and grown to ∼2⨯10^8^ CFU/ml. Growth curves were established by CFU at indicated times after the addition of stresses (0.3U/L xanthine oxidase, 1mM DEA NO/NOate, 2.5mM H_2_O_2_, or 2.5 mM peroxynitrite).

### ATP assays

HG003 was grown to ∼2×10^8^ CFU/ml in MHB. Where indicated cells were exposed to 0.3U/L xanthine oxidase, 1mM DEA NO/NOate, 2.5mM H_2_O_2_, or 2.5 mM peroxynitrite. After 0.5h, ATP levels were measured in 100µl cells as described previously(12) using a BacTiter Glo kit (Promega) according to the manufacturer’s instructions. For ATP measured following internalization in macrophages, unstimulated and stimulated macrophages were infected as above for 4h, followed by lysis and resuspension of *S. aureus*. Uninfected controls of both stimulated and unstimulated macrophages were used for background normalization. Following resuspension in PBS, cells were centrifuged at 300g for 5min to collect cellular debris, then at 10,000xg for 3min to pellet bacteria. Supernatant was removed, cells were washed 1x in PBS, and resuspended in a final volume of 200ul PBS. ATP levels were measured in 100μl cells as above and normalized to CFU. Averages and standard deviations of 3 biological replicates are shown (n=3). Statistical significance was calculated using the Student’s t-test (unpaired, two-tailed) or One-Way ANOVA with Sidak’s multiple comparison test as described in the figure legends.

### Aconitase activity

HG003 or mutant strains were grown to mid-exponential phase in MHB and incubated with or without 2.5 mM peroxynitrite. After 0.5h, 2ml cells were pelleted and resuspended in 200μl assay buffer and lysed with 50µg/ml lysostaphin at 37°C for 5min. Samples were pelleted and the supernatant was assayed for aconitase activity using an Aconitase Assay Kit (Cayman Chemical) without activation as per the manufacturer’s instructions. Aconitase activity was normalized to CFU. Averages and standard deviations of 3 biological replicates are shown (n=3). Statistical significance was calculated using the One-Way ANOVA with Sidak’s multiple comparison test as described in the figure legends.

### Immunoprecipitation and Western Blot of AcnA-YNO_2_

Unstimulated and stimulated J774 macrophages were seeded at 6.5⨯10^5^ cells/ml in supplemented MEM in 10cm treated dishes and incubated overnight at 37°C, 5% CO_2_. *S. aureus* strain HG003 *acnA::erm* pEPSA5_*acnA*_FLAG was grown in MHB media for ∼20h. After 20h, cultures were induced with 0.2% xylose for 1.5h, pelleted and resuspended in PBS. Macrophages were infected at MOI 100. Plates were incubated for 30min at 37°C, 5% CO_2_. After 30 minutes, the cells were washed 2x with PBS and fresh media containing 30µg/ml gentamicin (Fisher) was added to kill extracellular bacteria(51). After 4h incubation, cells were washed 3x with PBS. 5ml 0.1% Triton X-100 was added to each well for 5min at 37°C to selectively lyse the macrophages and release the bacteria. PBS was added to each plate, lysates were suspended by pipetting, and transferred to conical tubes. Samples were centrifuged at 10,000xg for 1 min to pellet bacteria and resuspended in 500ul PBS. Bacterial cells were lysed with lysostaphin (10μg/ml) for 10min at 37°C, or until confluent lysis was observed. After lysis, 2x protein inhibitor complex (Sigma) in PBS was added and protein levels were normalized by Bradford assay. Samples were mixed with 75ul mouse anti-FLAG magnetic beads (Sigma) and incubated at 4°C for 1h. After 1h, beads were washed 4x for 5min each with 1x PIC in PBS. Samples were boiled in SDS-reducing buffer and run on a 4–12% bis-tris acrylamide gel (Invitrogen). Protein was transferred onto a PVDF membrane, and YNO_2_ was detected using mouse monoclonal anti-YNO_2_ antibodies (1:500; Santa Cruz Biotechnology) with goat anti-mouse IgG-HRP secondary (1:10000; Cayman Chemical). Western blot was analyzed and quantified by densitometry using FIJI.

### Statistical information

Statistical method and sample size (n) are indicated in the methods for each experiment. Statistical analysis was performed using Excel (Microsoft) or Prism 8 (GraphPad) software.

## Acknowledgments

This work was supported by the NIH grants R01AI137273 to B.P.C., R03AI148822 to S.E.R. and Burroughs Wellcome Fund Investigators in the Pathogenesis of Infectious Disease (BWF-PATH) award to B.P.C. We are grateful to Jeffrey Boyd for providing strains. We are grateful to Janelle Arthur for equipment.

## Author Contributions

B.P.C, S.E.R. N.J.W, E.S.M.B, and J.E.B conceptualized the project; B.P.C, S.E.R., and J.E.B wrote the manuscript; V.G.F provided resources; J.E.B and J.C.S performed the in vitro experiments; N.J.W. and J.E.B performed the tissue culture experiments; S.E.R., N.J.W. and J.E.B produced figures; B.P.C and S.E.R provided funding for the project.

## Data availability

Additional data that support the findings of this study are available from the corresponding author, Brian P. Conlon, upon request.

